# An algorithm-based investigation reveals the differential dynamics of water inside protein cavity as a function of distance from its wall

**DOI:** 10.1101/2024.11.06.622213

**Authors:** Abel Xavier Francis, Mansi Chilkoti, Atul, Mrinal, Sukriti Sacher, Arjun Ray

## Abstract

Hydration forces exerted by water in the form of hydrogen bonding networks or electrostatic interactions play an essential role in protein structure and function. These interactions often govern chemical catalysis, ion transport, protein stability, and folding. While water’s role as a biological solvent and on the protein surface is widely studied, its function inside protein cavities is often neglected due to the existing challenges in its detection using experimental and computational approaches. The importance of studying these special protein-water interactions is further underscored by the fact that water spatially confined within cavities exhibits deviations from bulk behavior, directly impacting processes occurring inside protein cavities. With these challenges in mind and building upon our method that accurately identifies the protein inner cavity surface (CICLOP), we have developed a tool that can accurately distinguish water occurring within cavities from the bulk solvent around the protein. Our tool can characterize the dynamic properties of water within protein cavities, such as diffusion, residence time, and rotational and orientational relaxation, using molecular dynamics (MD) simulation trajectories as input. We demonstrate the robustness of our tool on several cavity-containing proteins and describe its applicability in characterizing the biological function of water confined within the cavity of an archaeal group II chaperonin.

## Introduction

Discussions of biological processes are incomplete without assessing the role of water. Indeed, water is central to life; cells, organisms, and entire ecosystems are completely dependent on the presence of water. Chemically, water is a polar, protic solvent that can ionize both itself and other molecules. This property enables it to form large networks of hydrogen bonds that form hydration layers with variable thickness around biomolecules[1]. This “biological water” is adaptable to changes in biomolecular conformations [2] and enables several biological processes, including protein folding [3], protein conformational changes [3], ligand binding [4] and twisting of DNA double helices [5] to name a few. Water molecules can also exist within the surface cavities, clefts, tunnels, channels, and within internal cavities in proteins. It is argued that even large hydrophobic cavities in proteins are also stabilized by clusters water molecules inside them [6].

Due to the close proximity with the functional groups on the surface of proteins and in the cavity lining interface, water in the first hydration shell and that confined within cavities behaves differently than water located far away from the surface of the protein (bulk–like water) [7, 8, 1]. The number density of these water molecules (governing the thickness of hydration) and their dynamic properties like rotational and orientational states, residence time, diffusion, and exchange kinetics are different from bulk-like water [9, 10, 11, 12, 13, 14]. These changes in the dynamic properties of water often drive biological processes. For instance, in small cavities of protein kinases, water molecules located in the catalytic pocket have high residence times (≈ 27.8*ps*) and were observed to stabilize important catalytic residues in several protein kinases [15]. Similarly, individual water molecules in the S1 pocket of trypsin-like serine proteases have been shown to drastically alter ligand binding affinity on complex formation [4]. In long cylindrical cavities of channel proteins having radii between 3 − 6 Å, the rotational reorientation of water inside the cavity was observed to be reduced in comparison to bulk water, altering the local dielectric constant within the pore, affecting its ion permittivity [9]. While in large internal cavities such as those in molecular chaperones, the water density inside the cavity has been proposed as an important parameter for efficient protein folding [16, 17].

Despite the fundamental role of “confined water” within protein cavities, determining the exact number of water molecules in cavities and measurements of their dynamics remains challenging by traditional experimental techniques. Both small angle X-ray scattering (SAXS) and neutron scattering (SANS) have been used to detect water in the interior of the protein, but these techniques do not provide any information about the exchange dynamics of this water. Furthermore, due to the positional disorder and low occupancy, especially within hydrophobic cavities, the accuracy of detection is limited [18, 19]. Nuclear magnetic resonance (NMR) and infrared spectroscopy have also been used to study the reorientation times of individual water molecules confined within cavities; however, the orientational disorder and shorter residence times of this water may obscure accurate measurements [18]. Femtosecond dynamic solvation shift (DSS) assays have also been widely used to study solvation dynamics, however concerns regarding the choice of probe and its effect on the assay has limited their use in case of proteins [20]. Similarly, techniques like dielectric relaxation (DR), depolarised light scattering (DLS), optical Kerr-effect (OKE), and terahertz (THz) spectroscopies have also been used to characterize collective relaxation or reorientational dynamics of water with the ability to detect slowdown/speedup in dynamics, however, deconvoluting these spectra in context of confined water is very challenging, requiring assumptions about the underlying relaxation time distributions.

On the other hand, computational approaches using molecular dynamics (MD) simulations have shown promise [9, 21, 18]. Significant advancements in force-field parameterization have led to accurate water models that can reproduce experimentally determined macroscopic properties of water, such as density, coordination number, surface tension, and more [22, 23]. Additionally, the spatial and temporal resolution provided by MD trajectories allows one to determine both the position and the dynamical properties of water in hydration layers and within the protein cavities [8]. While the vast array of work done in this area has focused on the dynamics of macromolecule hydration, very few studies have looked at the properties of water confined within the nanoscopic cavities of proteins[9, 10], with the majority of studies focusing on smaller cavities [18, 19, 24][24]. Water within such biological confines, unlike water in the putative hydration layers, has no direct contact with bulk water and hence cannot exchange with the bulk solvent at the same time scales.

The challenge lies in being able to distinguish the bulk solvent from the confined water. Previously, a classification scheme based on the coordination sphere of water molecules was proposed to distinguish bulk water (that has five or larger coordination numbers) from internal or surface water (that has zero to three water molecules in the vicinity) [25]. While extremely useful for small cavities, this classification scheme was observed to perform poorly on large internal cavities that may contain sizable water aggregates. Subsequently, the focus shifted to identifying internal cavities and the water within its vicinity [26]. Identification of protein cavity is an old problem with several methods addressing it using either a geometrical, probe-based, or tessellation-based approach [26, 27]. But neither of these methods inherently detects the water within the cavity. Moreover, a vast majority of these methods are limited by their accuracy, automation, and comprehensiveness in characterizing the cavity itself, impacting the inferences made with respect to its hydration.

To address these challenges, we have expanded our tool for the Characterization of Inner Cavity Lining of Proteins (CICLOP) to now detect the water within the protein cavity in addition to the cavity lining residues. Additionally, the method can take any MD simulation trajectory (independent of the MD simulation suite used) and characterize the dynamical properties of water identified within the cavity, such as its residence time (i.e., the time spent by a water molecule in the cavity before mixing with the bulk), diffusion and rotational relaxation. Due to CICLOP’s robust and fast algorithm for cavity detection at an atomistic resolution [27], an accurate identification of water around the cavity is achieved such that not a single water molecule in the cavity is missed. Our method is available as a standalone, allowing users to perform in-depth characterization of water dynamics within protein cavities with its easy-to-use modules. In this work, the algorithm for cavity water identification, along with the different modules for water dynamics characterization, are discussed. Our method’s applicability is also presented by characterizing the water within the internal cavity of *Methanococcus maripaludis*, Mm-Cpn, an archaeal group II chaperonin that is involved in protein folding. Using MD simulation of the closed states of Mm-Cpn, we show that the dynamics of water within the protein folding chamber is slowed down and how it may impact protein folding.

## Results

### Overview of our computational framework to identify water within the cavity

This method serves as an extension of our inner cavity identification tool, CICLOP, which identifies residues lining the protein cavity. Instead of taking a single PDB file as input, we now take molecular dynamics trajectories as input and identify the inner surface of the protein in each frame of the trajectory. Using this inner surface as a reference, all the oxygen atoms of water at varying distances from the protein inner surface boundary in each frame of the trajectory are marked and stored as cavity water **(Sup Fig 1)**. For every water molecule, based on the position of the oxygen in the voxel grid of 1Å^3^, its position with respect to the cavity’s inner lining is recorded. A B-factor loaded PDB is provided as an output where the residues/atoms lining the cavity are marked with a B-factor of 9999, water within 10 Å from the cavity lining is marked with a B-factor of 1000, while water within 10 − 20 Å and beyond 20 Å is marked with a B-factor of 4000 and 7000 respectively. This B-factor loaded PDB can be easily visualized using any structure visualization software to render publication-quality images depicting the water inside the protein cavity **(Fig 1)**.

**Fig 1:**
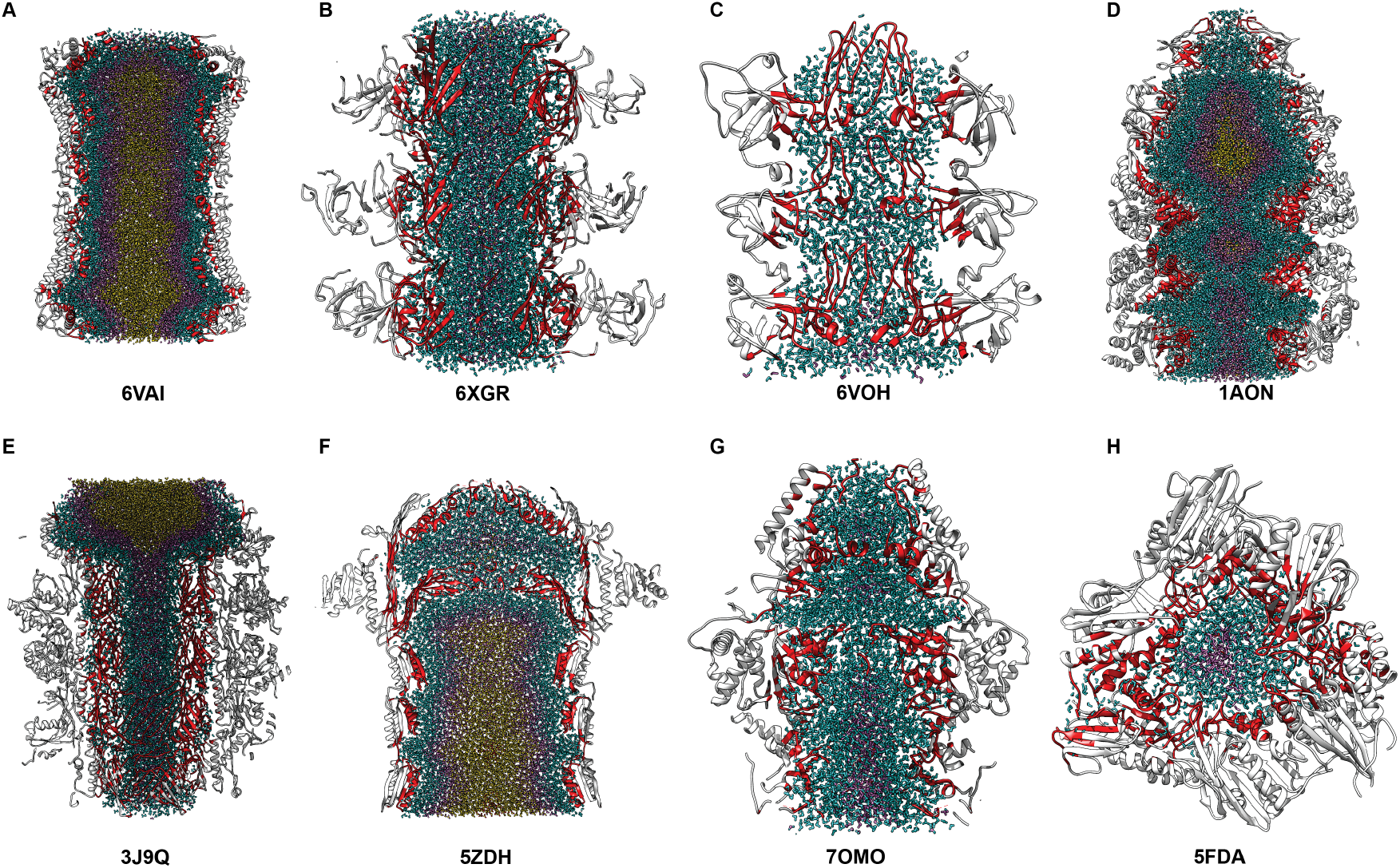
Visualizing water within protein cavities. Identification of residues lining the inner cavity and water in its vicinity using our algorithm for **A**. human calcium homeostasis modulators (CALHMs) **B**. tail tube protein of bacterial flagella **C**. ATP synthase of chloroplast **D**. GROEL/GROES chaperonin complex **E**. bactericidal contractile structural protein **F**. bacterial pilotin-secretin AspS-GspD complex **G**. Renilla reniformis luciferase **H**. apo form of dihydrofolate reductase. Protein residues are depicted in white, while cavity lining residues are shown in red. Water within 10 Å of the cavity lining is shown in blue, while water within 10 − 20 Å and water beyond 20 Å is shown in purple and yellow, respectively.

Our mixed approach of using grids and tessellations to identify inner residues offers a significant advantage over the probe-based (sphere) approach, as it can delineate the protein’s inner surface with precision. As a result, not a single water molecule in the cavity (irrespective of its position, whether close to the protein boundary or deep within the cavity) is missed. Furthermore, our algorithm is not dependent on any MD simulation suite and can be easily scaled up to include new functionalities in the future.

We have tested this method on a large subset of cavity-containing proteins varying in size, shape, and function. We can efficiently detect water within hydrophobic and hydrophilic cavities that are either curved or disconnected. **Fig 1** showcases the water within the internal cavities of proteins with diverse cavity architectures, where the cavity residues detected by our tool are colored in red and water within 10 Å, water between 10 − 20 Å, and water beyond 20 Å are depicted in blue, purple and yellow respectively. The accuracy of our method is evident by the distinct boundaries of water observed in each of these cavities. Alternatively, water within smaller cavities like that in **Fig 1H** is also detectable, and that in large internal cavities with chambers formed due to fenestrations in the cavity is also clearly distinguishable **(Fig 1C-G)**.

In addition to the identification of cavity lining (at the residue and atomic level) and the confined water, our method provides an exhaustive package of easy-to-use features and modules that can characterize the cavity and the dynamics of water within the cavity. All the previous functionalities of CICLOP have been retained, such that the method still characterizes the cavity lining residues based on their conservation scores and secondary structure. Instead of the diameter and radius profile for a single PDB structure, our method now generates the cavity dimension profiles for every frame in the trajectory, providing the user with an average dimension of the cavity throughout the simulation with error bars that depict the fluctuations in dimensions from the average. A charge profile of the cavity generated from the localization of charges along the inner cavity surface is also calculated. However, instead of adding up the charged residues that line the cavity, partial charges on each atom that is found to be a part of the inner lining are summed up along the protein axis, making the new charge profile much more accurate than the one generated by CICLOP previously **(Sup Fig 2)**.

For characterizing the cavity water, our method computes the cavity water density profile in layers of 1 Å from the cavity lining towards the cavity center **(Sup Fig 3)**. Furthermore, the user has the option of choosing the region within the cavity and the duration within their simulation for which they intend to calculate other useful dynamical properties of water. The most important physical variable being, the residence time of all the water within the cavity or of the water within certain regions of the cavity, can be calculated efficiently using our method. Furthermore, using these residence times as a reference, the dipole rotational relaxation time of water in the entire cavity or confined in certain regions of the cavity can also be calculated. Similarly, diffusion dynamics can also be calculated. Our module also enables the user to study the propensity of movement of water within the cavity.

Due to the extremely fast dynamics of water, it is advisable to use MD simulation trajectories with an integration time step of 2 *fs* or smaller with a maximum saving frequency of 100 *fs*, such that the water molecules are efficiently tracked during the dynamics analysis.

### Characterizing the dynamics of water within the Mm-Cpn cavity using our method

To demonstrate the usage and applicability of our method, we used our method to study the water dynamics within the cavity of an archaeal thermosome chaperonin from `Methanococcus maripaludis` (Mm-Cpn) PDB ID: 3LOS. Mm-Cpn assists in the folding of nascent and misfolded proteins in the cytosol of archaebacteria; however, the manner in which it recognizes and interacts with unfolded intermediates is incompletely understood. Structurally, it consists of two rings with eight subunits each, together forming a 16-subunit closed homo-oligomer that has a large internal cavity at its center. The length of the entire cavity end-to-end is 150 Å, while the diameter of two hemi-spheric chambers formed are 160 Å each and 104 Å at the region where they come together in the center **(Sup Fig 4)**. This trend is also reflected in the volume profile of the cavity **(Fig 2E). Fig 2A & B** depict the vertical and transverse sections of the protein, highlighting the large internal cavity enclosed within the chambers. The residues lining the cavity are colored in red.

**Fig 2:**
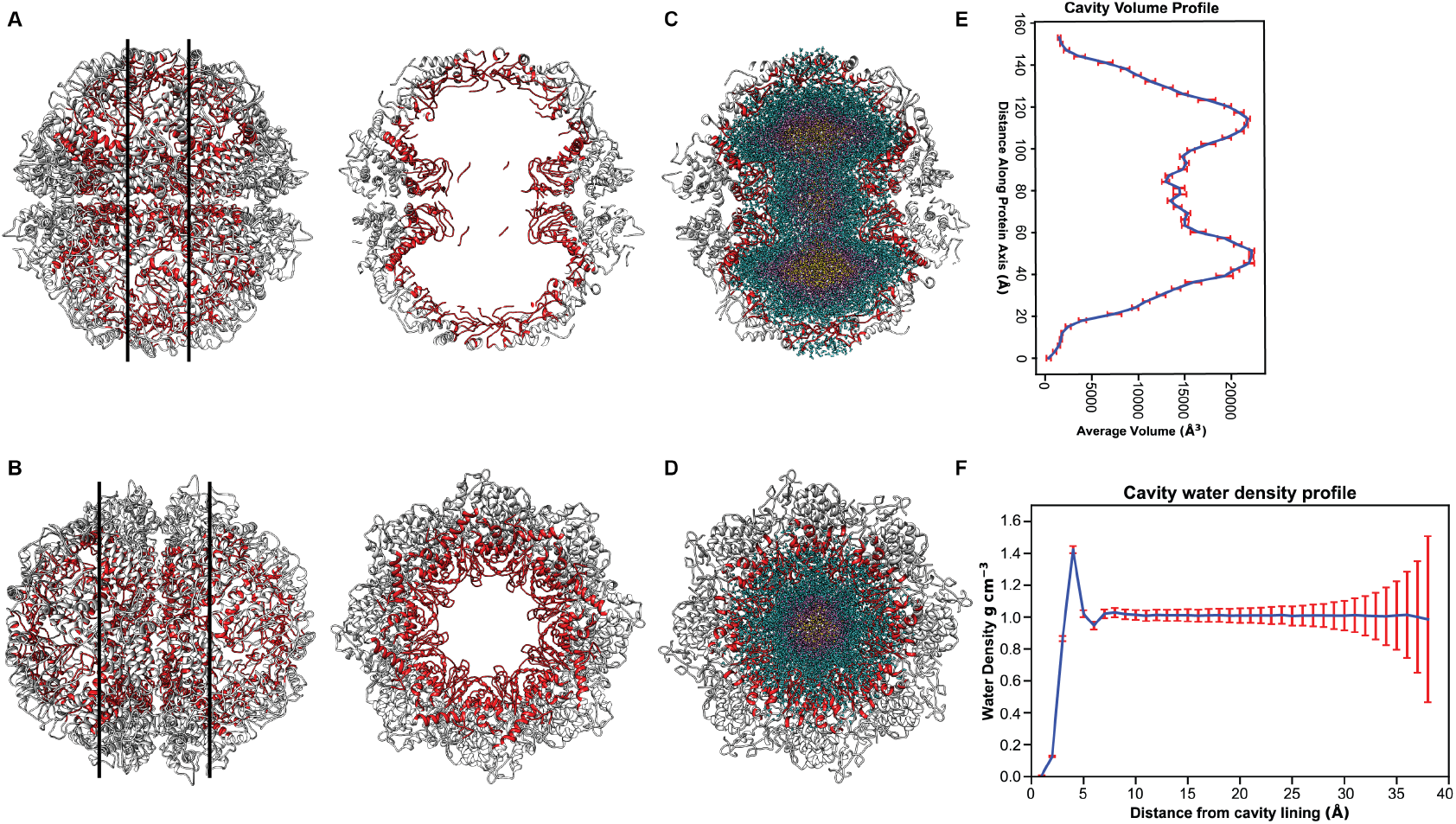
Characterizing water dynamics of the closed state of chaperonin from Methanococcus maripaludis (Mm-Cpn). **A**. Transverse section **B**. Horizontal section of Mm-Cpn cavity highlighting the cavity dimension. Protein residues are depicted in white, while residues lining the cavity are shown in red. Hydrated **C**. transverse and **D**. horizontal section of the cavity, such that water within 10 Å of the cavity lining is shown in blue, while water within 10 − 20 Å and water beyond 20 Å is shown in purple and yellow, respectively. **E**. The volume profile of the inner cavity along the protein axis when it is aligned to the *Z*-axis. **F**. Water density profile with distances computed from the cavity inner lining moving towards the center of the cavity.

Following the MD simulation of the closed state of Mm-Cpn **(Sup Fig 5)**, we identified the water within the cavity throughout the simulation and calculated its density using our improved method. The density profile shows an increased density (1.4 *g cm*^*−*3^) of water within 5Å from the cavity wall, beyond which it plateaus to the bulk water density of 1 *gcm*^*−*3^ **(Fig 2F)**. This profile clearly highlights the clustering of cavity water closer to the cavity wall.

To understand this behavior of water, we calculated the charge profile of the cavity by summing up all the partial charges of atoms that line the inner surface lining. We observed that the cavity is predominantly negatively charged, with the total cavity charge between −179*e* and −121*e*. Interestingly, the maximum magnitudes of negative charges correspond to the regions of the cavity having the largest radii **(Sup Fig 2 & 4)**, which are proposed to be the folding chambers of Mm-Cpn. Furthermore, the increased density of water molecules around the cavity wall may therefore be due to an increased H-bond propensity of water in this region with the negatively charged atoms lining the cavity interface.

### Water closest to the cavity wall in Mm-Cpn has high residence times

To study the dynamical properties of water, such as translational diffusion coefficients and rotational relaxation times, it must be ensured that the ensembles of water under consideration, on average, stay within a predefined area for the time period over which the analysis is being done. Therefore, the residence times of the water within the cavity were calculated. The residence time of total water (*τ*_*Res*_) within the cavity was observed to be 23.07 *ps* **(Sup Fig 6)**. Next, we looked at residence time of water at varying distances from the cavity wall (within blocks of 5 Å each) moving toward the center of the cavity. We observed that the *τ*_*Res*_ of the water closest to the cavity wall (≤ 5 Å) is 7.13 *ps* **(Fig 3A)**. However, for water between 6 − 10 Å from the cavity wall, *τ*_*Res*_ is 0.85 *ps* **(Fig 3B)**, which continues to decline to 0.71 *ps* and 0.67 *ps* for the next 5 Å blocks, respectively **(Fig 3C & D)**. Looking at the residence time values and the density plot, together with the charge profile of the cavity, it is evident that for the chosen Mm-Cpn system, water prefers to stay near the cavity wall because it is negatively charged.

**Fig 3:**
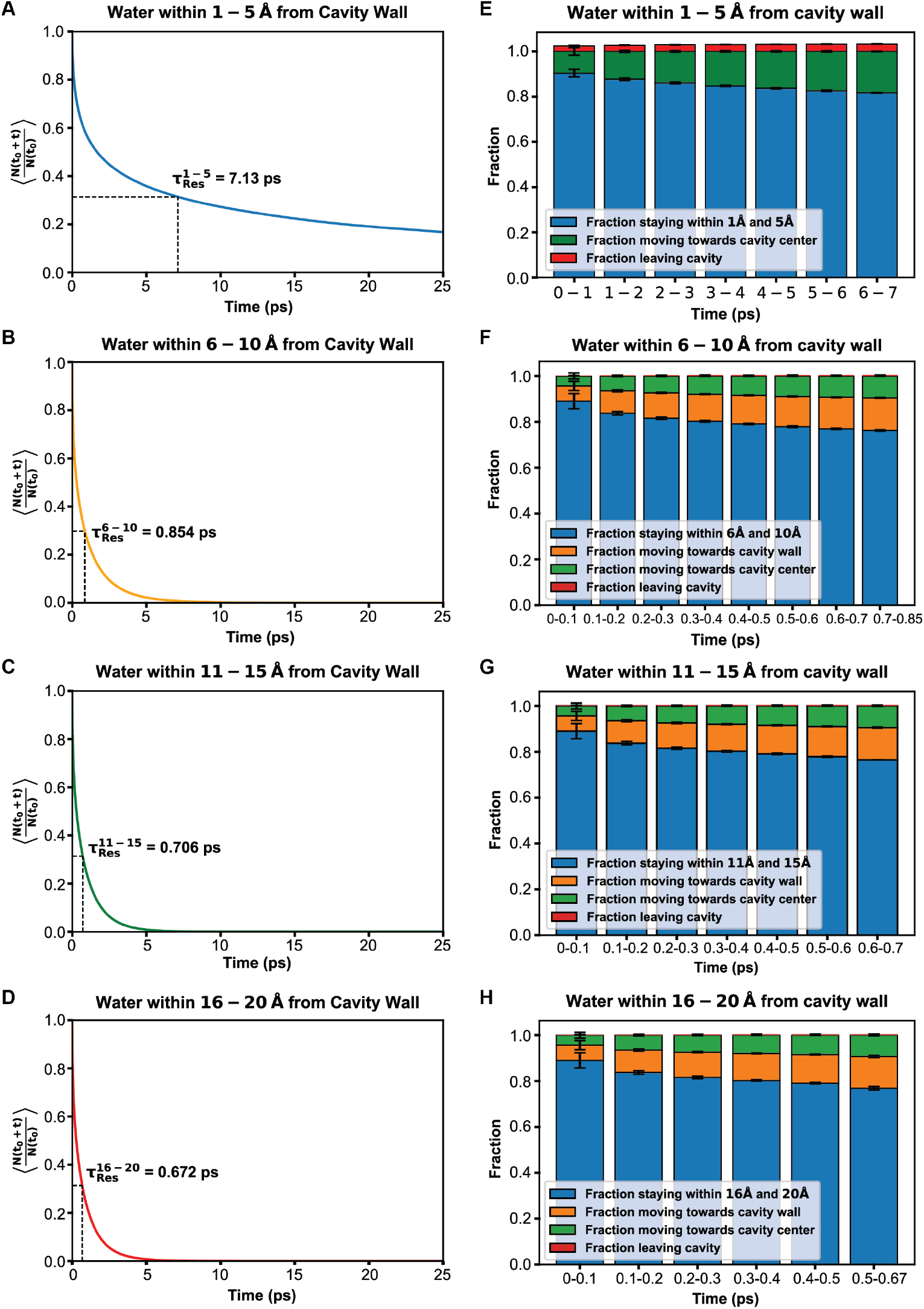
Characterizing the water dynamics within the regions of the MmCpn cavity. Residence times of water within **A**. 1 − 5 Å **B**. 5 − 10 Å **C**. 10 − 15 Å **D**. 16 − 20 Å from the cavity wall. Propensity of water within **E**. 1 − 5 Å **F**. 5 − 10 Å **G**. 10 − 15 Å **H**. 16 − 20 Å regions of the cavity to move to other regions within the cavity.

Next, based on the residence times of water, we evaluated if the water preferentially moves to a certain region of the cavity. Using the inbuilt water movement propensity module in our method, we identified the number of water molecules in the cavity at *t*_0_ and then counted the number of water molecules that stay within layers of 5 Å each from the cavity wall after some time (*t*_0_ + *dt*), thus giving a general idea of the exchange kinetics between the layers. For layers closest to the cavity (≤ 5 Å), we observed that 80 − 85% of the water stayed near the cavity wall, while 10 − 15% of water moved towards the center of the cavity. Only a very small fraction of water (3%) entirely left the cavity **(Fig 3E)**. For all the other layers, 80 − 85% of water was observed to stay within the layer, while 10 − 15% was observed to move towards the cavity wall **(Fig 3F-H)**. This preferential movement towards the cavity wall may have resulted in an increased density of water molecules close to the cavity wall observed in **(Fig 2F)**.

### Water closest to the cavity wall in Mm-Cpn has very slow diffusion

We then calculated the translational diffusion coefficient of the total water within the cavity and of the water ensembles based on their distance from the cavity wall within their respective residence times. The value of diffusion coefficient (*D*) for total water within the cavity was observed to be 3.38 × 10^*−*5^ *cm*^2^ *s*^*−*1^ **(Sup Fig 7)**. While for the water within ≤ 5 Å from the cavity wall was observed to be 1.514 × 10^*−*5^ *cm*^2^ *s*^*−*1^ **(Fig 4A)**, while for water between 6 ≤ 10 Å from the cavity wall, *D* = 4.618 × 10^*−*5^ *cm*^2^ *s*^*−*1^ **(Fig 4B)**. For layers between 11 ≤ 15 Å and 16 ≤ 20 Å from the cavity wall, the diffusion coefficient was observed to be 6.044 × 10^*−*5^ *cm*^2^ *s*^*−*1^ and 6.445 × 10^*−*5^ *cm*^2^ *s*^*−*1^, respectively **(Fig 4C & D)**. We similarly calculated the diffusion of bulk water for a system of 4055 TIP3P water molecules, *D*^*Bulk*^ = 5.49 × 10^*−*5^ *cm*^2^ *s*^*−*1^ **(Sup Fig 7)**.

**Fig 4:**
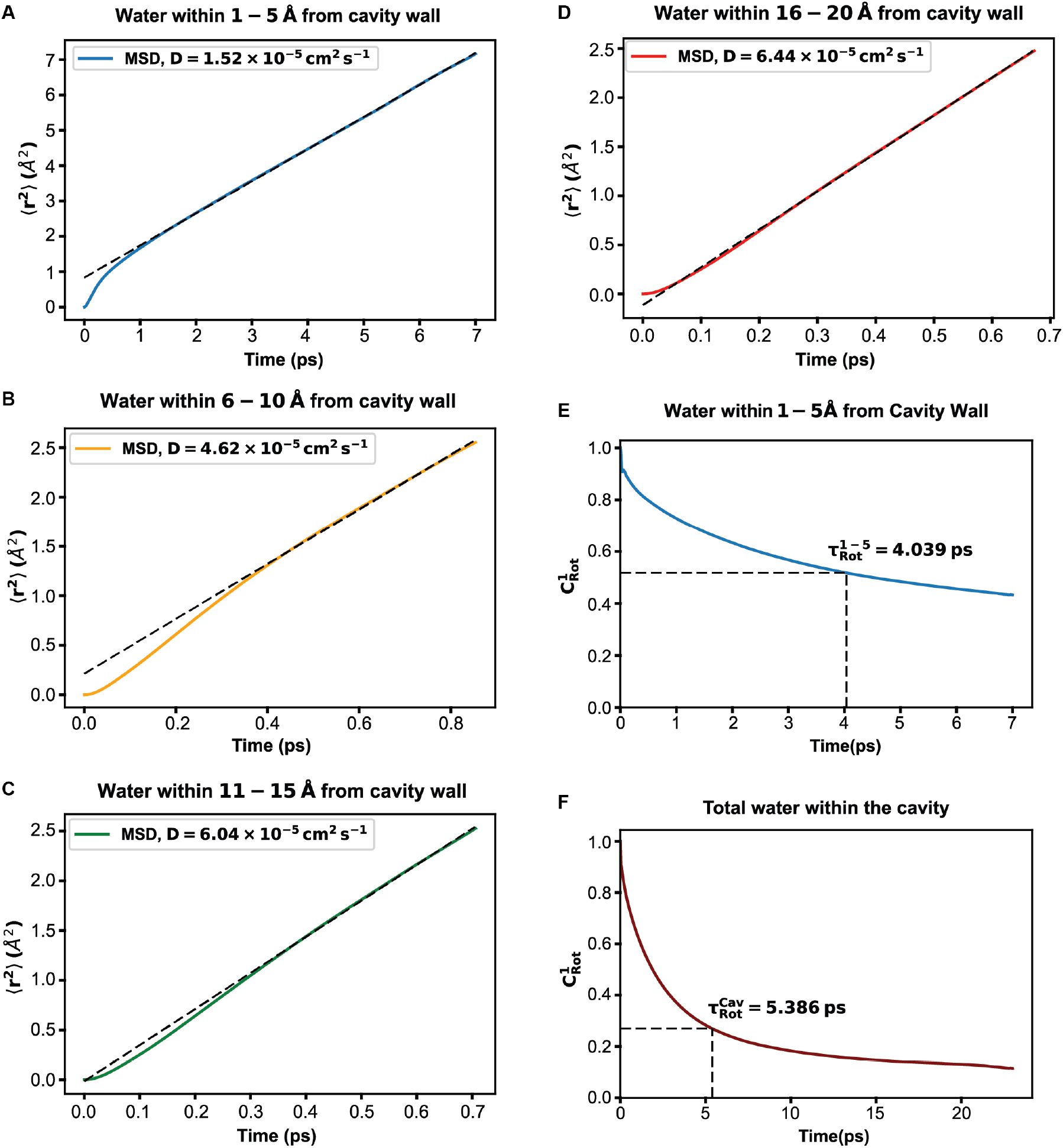
Diffusion and rotational relaxation of cavity water. Diffusion profiles showing the diffusion coefficient of water within **A**. 1 − 5 Å **B**. 5 − 10 Å **C**. 10 − 15 Å **D**. 16 − 20 Å from the cavity wall. Rotational relaxation of water within **E**. 1 − 5 Å and **F**. total water within the cavity.

We observed that the water closest to the cavity wall undergoes slow diffusion (3.6 times less than the bulk water), while water farther from the cavity wall approaches *D*^*Bulk*^. It must be noted that for water beyond 10 Å from the cavity wall, the mean square diffusion coefficient is calculated for a very short period of time (*<* 1*ps*), as the residence time of water in these regions is very less. Therefore, it is much more likely that the *D* value obtained is from the ballistic section of the MSD curve and, hence, higher than *D*^*Bulk*^ itself. Interestingly, water in protein hydration shells have also been shown to exhibit 2-3 times slower diffusion dynamics than the bulk water [8, 28]. This modest but distinguishable decrease must be a result of the solute perturbation.

To further characterize the degree of perturbation of cavity lining residues on water within the cavity, we calculated the rotational relaxation of total water within the cavity and water within ≤ 5 Å from the cavity wall. The rotational reorientation time of water molecules within the cavity represents the amount of time taken on average for the dipole vector of a water molecule to get de-correlated from its initial configuration. For the water closest to the cavity wall, the rotational relaxation time was observed to be 4.309 *ps*, while that of total cavity water was 5.386 *ps* **(Fig 4E & F)**. This, in comparison with bulk water 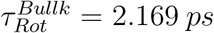 **(Sup Fig 8)**, implies that the water within the cavity has a slower rotational relaxation by a factor 2, highlighting the sluggish dynamics of water within the folding chambers of Mm-Cpn.

## Discussion

We have developed a new Python-based package for the identification and characterization of water and its dynamics in protein cavities. We have focused on providing easy-to-use functionalities for water dynamics characterization, such as water density, residence time, rotational relaxation, and diffusion kinetics. This package can be used for the characterization of static protein structures or molecular dynamics simulation trajectories obtained from any MD simulation suite. This tool, available as an executable, is fully unit-tested and has extensive documentation for its usage. The current implementation uses MDAnalysis [29] for atom selection and trajectory reading. This enables us to make use of the powerful MDAnalyis atom selection language for selecting specific regions of the cavity, ensuring that our method can be used for non-protein systems as well. Additionally, its modularity and multi-processing capabilities allow it to extract meaningful information about water dynamics from trajectories spanning longer time scales.

We have also provided an example of our method’s use in probing the cavity water dynamics of a group II chaperonin from Methanococcus maripaludis (Mm-Cpn). Group II chaperones are important protein-folding machines found in organisms ranging from archaea to humans and have been actively studied. It has been proposed that following the substrate binding, an ATP-mediated conformation change leads to the closure of folding chambers in Mm-Cpn [30, 31, 32]. Further cryo-EM studies have revealed that upon the closure of folding chambers, the substrate is released into the cavity of the chaperonin[33], where it can undergo folding protected from the cytosolic environment inhibiting the formation of non-native structures, or toxic aggregates. Therefore, the nature of the environment within the cavity, especially the water confined within it, is likely to play an important role in the folding of the confined substrate, as has also been proposed for the GroEL/ES chaperonin complex[34, 35, 36].

Our observations have shown that water within the Mm-Cpn cavity exhibits drastically slower dynamics as compared to bulk water, with water closest to the cavity wall exhibiting the largest deviations. When a substrate is bound within the Mm-Cpn cavity, it will likely occupy the central regions of the cavity, displacing the water within those regions. Since the folding chambers are closed on substrate binding, the probability of water moving out of this cavity is significantly reduced. Additionally, our observation shows that water within the central regions of the cavity has an increased propensity to move toward the cavity wall since it is significantly negatively charged. This exclusion of water away from the bound substrate and towards the cavity wall may lead to a diminished hydrophobic effect within the central regions of the cavity as proposed by Koroboko *et. al* [34] in the case of the GroEL/ES chaperonin complex. This diminished hydrophobic effect would weaken hydrophobic contacts, enabling the refolding of aggregates or misfolded substrates within the Mm-Cpn cavity. Hence, the altered dynamics of water within the Mm-Cpn cavity may be responsible for producing a pro-folding environment for protein substrates.

## Conclusion

With our method, it is now possible to characterize cavities and the dynamics of water within these cavities. Since these cavities are isolated from the bulk, the water within the cavity does not freely exchange with the bulk solvent. This distinguishes cavity water from the protein hydration layer that coats the outer surface of the protein. Therefore, the theoretical approaches previously undertaken to study protein hydration that assume a free equilibrium between the bulk solvent and the hydration layer [13] can not be directly applied to cavity-confined water. However, the diffusion and rotational orientation of water within this exemplar cavity of Mm-Cpn is observed to be 2-3 times slower than the bulk water, as is the case with water in hydration layers [8]. This reinstates the intensity of perturbations caused by chemical groups lining the cavity and on the surface of the protein. Regardless, the change in dynamics of water closest to the perturbing groups can influence biological function.

We believe that the versatility of our tool and its ability to accurately and robustly characterize protein inner cavities and the water dynamics within these cavities will have a significant application in the field of structural biology in evaluating protein structures, protein oligomerization studies, substrate binding, functional characterization of channels and in understanding chaperone-assisted protein folding. Further, its ability to identify not just water, but any molecule within protein cavities would be invaluable for the study of small molecules, ions, lipids and other biomolecular entities confined within protein cavities. We may be limited to just identifying these entities within the cavity for now, but in the future, we intend to add more modules capable of tracking and characterizing their dynamics. We also intend to incorporate modules for the calculations of more dynamical and thermodynamic variables of water within protein cavities.

## Materials and Methods

The current method for cavity water detection and characterization is an expansion of our previously developed tool for the characterization and identification of protein inner residues (CICLOP)[27], which can now detect the cavity lining and the water within the cavity. The basic algorithm for the identification of cavity residues is the same and is briefly described here for continuity.

### Algorithmic overview for cavity detection: CICLOP

CICLOP takes a protein structure file as an input (PDB format only) and considers only the protein atoms. First, it obtains an initial approximation of the cavity axis by drawing a best-fit line to minimize the sum of the squares of the perpendicular distance of *Cα* atoms of the protein. This aligned protein is then mapped to a threedimensional grid of voxel cubes, where each voxel cube has a volume of 1 Å^3^. These voxels are bound by other voxels on each of their six sides, forming a connected network. Additionally, they may either be empty or contain one or more protein atoms. The voxels containing no atoms are marked empty, and for each of these empty nodes, adjacent nodes are searched to determine their occupancy using a breadth-first search. A cluster of empty nodes (voxels) may represent a cavity within the protein or the empty space beyond the outer surface of the protein; however, all of these empty nodes are recorded.

Next, the entire protein cavity is traversed along the protein axis in thin slices of 1 Å each. For each such slice

1. The geometrical center(*X*_*Cen*_, *Y*_*Cen*_) of all the atoms found to be in the boundary surfaces for that *Z* slice is found as

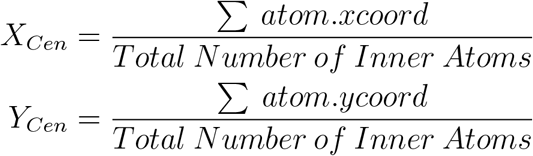
2. The mean radius(*R*_*Mean*_) at which the boundary atoms lie from (*X*_*Cen*_, *Y*_*Cen*_) is calculated.

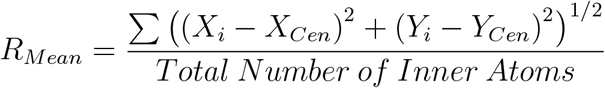
3. For every *Z* slice, the standard deviation of all the inner atoms which lie at a distance *R*_*i*_ ≤ *R*_*Mean*_ is calculated

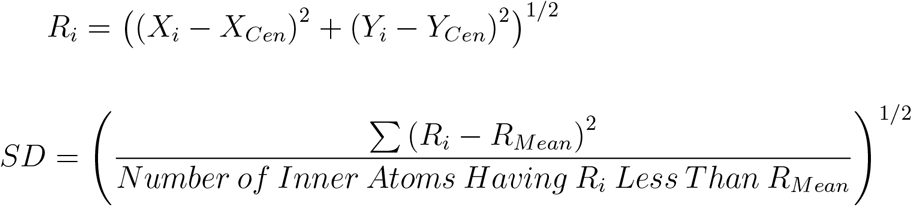
4. Now, every atom that lies outside a radius defined as *R* = *R*_*Mean*_ −0.7 × *SD* is unmarked as an inner atom and is discarded.
5. The above procedure is repeated for every 1Å − *Z* Slice and is concatenated to obtain all of the inner cavity lining atoms/residues along the axis of the protein.

Therefore, using the geometrical center and the mean radius (determined by the atoms found on the boundary surface: occupied voxels adjacent to empty voxels), a cutoff is calculated. Based on this cutoff, the outer protein surface is distinguished from the inner surface of the cavity. The algorithm continues to check for empty voxel nodes that may have been left unvisited to determine multiple cavities, if any, in the protein.

### Water detection inside the cavity

Once the inner surface atoms are identified, all the voxels that are empty and within *R* = *R*_*Mean*_ −0.7 × *SD* are identified as inner cavity voxels for every slice along the protein axis. Next, all the water molecules within these empty voxels are identified by mapping each oxygen atom of a water molecule to a voxel in the voxel grid.

To proceed further, all of the empty voxels within the cavity are assigned an attribute *layer value*, which defines the distance from the cavity wall that a particular voxel belongs to, allowing for the characterization of cavity water at varying distances from the cavity lining. The assignment of layer values to cavity voxels is done by first identifying the voxels within the cavity that lie just adjacent to the cavity lining atoms. These voxels are assigned a layer value of 1. Now, iteratively, all other voxels are assigned a layer value of *i* if it is found to be adjacent to another voxel having a layer value of *i* − 1. This produces natural bins of size 1 Å starting from the cavity lining toward the cavity center with the *i*^*′*^*th* layer containing water molecules lying at a distance of *i* − 1 Å to *i* Å from the cavity wall.

Since water identification within the cavity is dependent only on identifying the voxels within the cavity, we can also detect any molecule that lies within the cavity. A specialized input using the MDAnalysis selection language can be used to select any molecule of one’s interest to be searched for within the cavity.

Subsequently, a PDB file is written out for visualization containing all the protein and water atoms, such that the residues lining the inner surface of the cavity are marked with a B-factor of 9999, while the solvent molecules within 10Å from the cavity lining are marked with a B-factor of 1000, solvent molecules within 10-20Å from the cavity lining are marked with the B-factor of 4000 and those beyond 20Å are marked with a B-factor of 8000.

In order to characterize cavity and cavity water hydration dynamics, our method can now take MD simulation trajectories as an input (irrespective of the MD simulation suite used). The above algorithm is then applied to every frame in the trajectory to obtain time series data.

### Charachterization of the cavity in real time

We have retained all the previous modules for characterizing the inner cavity, except that they can now be run on MD simulation trajectories instead of just single protein structure files (PDBs). All the modules except the conservation score generate an average profile with error bars depicting deviations from the mean. The conservation score module from CICLOP has been retained as is. Moreover, the hydrophobicity profile has been discontinued. While the charge profile along the cavity axis is computed by summing up the partial charges of all the atoms lying on the inner surface instead of the charged residues. The cavity dimensions (radius and volume) are determined as previously described in [27] to generate an average radius and volume profile of the cavity.

### Water Density Plots

The layer-wise densities of the solvent oxygen atoms are calculated by identifying the number of solvent oxygen atoms present in the voxels of a particular layer and then dividing by the total volume of that particular layer.

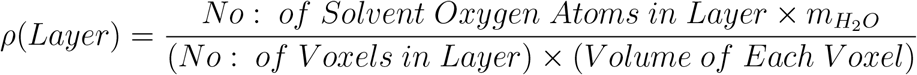

Where 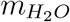 is the mass of a single water molecule in grams.

In a similar fashion, time-averaged plots for the oxygen number density are also generated. The layer-wise density information is generated for multiple frames in the simulated trajectory based on user inputs. It may be noted that since the dimensions of the cavity are not constant throughout the simulation, each frame in the trajectory contains a varying number of layers that describe the cavity. Therefore, for consistency, we perform the averaging based on the minimum number of layers detected in the cavity. The averages are also plotted along with their standard deviations as the error bars.

### Residence Time Calculations

The average residence time of water present at varying distances from the cavity lining can be calculated by selecting the water molecules between the bounding voxel layers *i* and *j*. We determine the survival probability of an ensemble of water as,

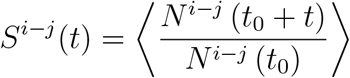

Where *N*^*i−j*^ (*t*_0_) represents the number of water molecules found between the layers *i* and *j*, at some initial time *t*_0_ and *N*^*i−j*^ (*t*_0_ + *t*) represents the number of initially identified water molecules that continuously remain between layers *i* and *j* after a time *t* has elapsed. The angular brackets represent an averaging over the time origins *t*_0_.

To obtain an average residence time, the above-obtained series of survival probabilities is integrated with respect to time. It must be noted that this method of calculating the residence time is model-independent and does not assume any particular form for the decay of the survival probability function.

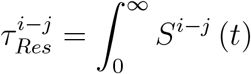

### Translational Diffusion Coefficients

The translational diffusion constant of a selected ensemble of water is calculated by using the well-known Einstein’s Diffusion Equation,

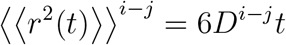

The double angular brackets correspond to an ensemble and a moving time average. The diffusion constant for the selected ensemble (*D*^*i−j*^) is found by fitting ⟪*r*^2^(*t*)⟫^*i−j*^ vs *t* to a straight line. While fitting, the initial and final 10% of the data is disregarded to prevent fitting to the ballistic domain of diffusion at lower time scales and to prevent fitting to regions at larger time steps where there is more statistical uncertainty due to comparatively worse time averaging. If one desires to change the window over which the fit is performed, the user can preemptively provide the start and end points over which the fit should be performed as user input.

### Rotational Relaxation Times

The rotational reorientation dynamics of water molecules within the cavity can be studied by calculating the First and Second Order Dipole Autocorrelation Function given by,

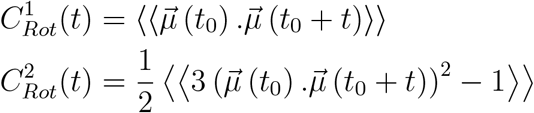

To obtain the Rotational Relaxation Time, we integrate 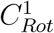 and 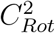 over the range of *t* over which it has been calculated.

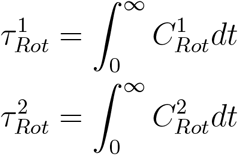

The user defines the ensemble of water under consideration by selecting an upper and lower bound for the voxel layers within the cavity and only the water molecules contained within these bounding voxel layers are considered for analysis.

### Water Movement Propensity

To identify whether the bias in the movement of water at a distance *d* from the cavity wall, defined by the bounding layers *i* and *j*, towards either the cavity wall, center of the cavity, or outside the cavity, we provide the functionality to calculate the movement of water from an initially identified ensemble over time.

From an initial time point *t*_0_, we calculate the following quantities, *F*_*W all*_ (*t*_0_ + *t*), *F*_*Center*_ (*t*_0_ + *t*), *F*_*Self*_ (*t*_0_ + *t*) and *F*_*Out*_ (*t*_0_ + *t*), the fractions of the initially identified water molecules moving towards the cavity wall, towards the center of the cavity, remaining in the initially defined region and leaving the cavity altogether after a time *t* has elapsed.

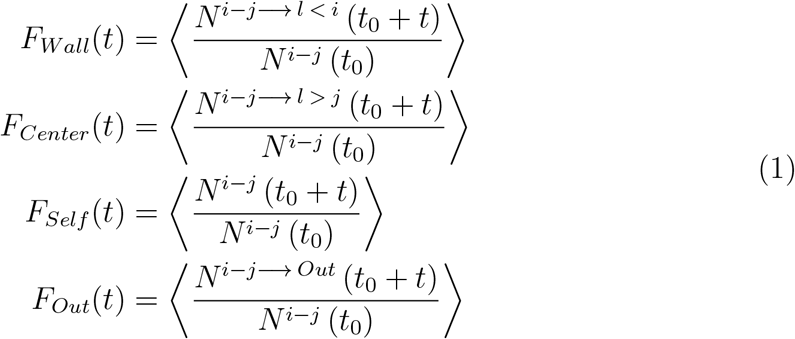

Here *N*^*i−j−→ l < i*^ (*t*_0_ + *t*) corresponds to the number of water molecules that have moved from the region bound by the layers *i* and *j* to some layer *l < i* in a time *t*. Since the assignment of layer numbers to voxels is directly representative of the proximity to the cavity wall, movement of water molecules to a layer *l < i* implies movement towards the cavity wall.

Similarly, *N*^*i−j−→ l > j*^ (*t*_0_ + *t*) represents the number of water molecules moving from the region bound by the layers *i* − *j* to some layer *l > j* in a time *t*, representative of movement towards the cavity center. *N*^*i−j*^ (*t*_0_ + *t*) represents the number of water molecules that remain in the same region after a time *t* and *N*^*i−j−→ Out*^ (*t*_0_ + *t*) represents the number of water molecules from the initial region that has moved out of the cavity after a time *t*. The angular brackets represent an average over time origins *t*_0_.

It is to be noted that the calculation of *F*_*Self*_ (*t*) differs from the calculation of the survival probability *S*^*i−j*^(*t*) in the fact that the calculation of the survival probability only counts those water molecules that have stayed continuously within the defined region for the entirety of the elapsed time *t. F*_*Self*_ (*t*) does not have this restriction and takes into account any and all water molecules that have left the defined region and returned in time *t*.

### Multiprocessing

Multiprocessing is a method of parallel execution where multiple processes run independently, allowing tasks to be split across multiple processing units or cores. This is especially useful for computationally expensive tasks that can be broken down into smaller, parallelizable components. Each process runs in its own memory space, which makes multiprocessing ideal for tasks where multiple frames or chunks of data need to be processed simultaneously without affecting one another. This approach maximizes CPU utilization, leading to significant performance improvements when analyzing large datasets

For the analysis of large structures with extensive trajectories, we leveraged Python’s multiprocessing libraries to optimize performance. In several instances, applying the same processes or functions across different frame steps within the trajectory was required. Given that these processes operate independently of one another across frames, parallel execution of the analysis on multiple frames was both feasible and efficient.

We implemented multiprocessing using the ***concurrent*.*futures*** module, which offers the `ProcessPoolExecutor` method. This method divides iterables into multiple chunks and submits them to the process pool as individual tasks. By default, the method splits the data into 10 chunks, although the user can customize the number of chunks based on specific requirements and the available computational resources. This flexibility allows for better resource management and enhanced performance for large-scale trajectory analysis.

We have also tested the time complexity of our algorithm and found it to increase linearly as system size or trajectory size increases **(Sup Fig 9)**.

### Molecular Dynamics Simulation of MM-Cpn

MD simulations were performed in triplicates with the GROMACS 2022.3 suite [37, 38, 39]. The system consisting of the closed structure of Mm-Cpn (**PDBID: 3LOS**) was represented by CHARMM36 [40, 41]. The water was modeled using TIP3P representation [42]. The starting conformations were placed in a dodecahedron box large enough to contain the system with at least 1.0 *nm* of solvent on all sides. Periodic boundary conditions were used, and the long-range electrostatic interactions were treated with the particle mesh Ewald method [43] using a grid spacing of 0.12 *nm* combined with a fourth-order B-spline interpolation to compute the potential and forces in-between grid points. The real space cutoff distance was set to 1.2 *nm* and the van der Waals cutoff to 1.2 *nm*. The bond lengths were fixed using the LINCS algorithm [44], and a time step of 2 *fs* for numerical integration of the equations of motion was used. Coordinates were saved every 10 *ps*. The pressure coupling was done by employing a Parrinello-Rahman barostat [45] using 1 *bar* as the reference pressure and a time constant of 2.0 *ps* with compressibility of 4.5 × 10^*−*5^ bar using the isotropic scaling scheme. Three hundred fifty-two positive counter-ions (Na+) were added by replacing an equal amount of water molecules to produce a neutral simulation box. All the starting structures were subjected to a minimization and equilibration protocol recommended by CHARMM. A simulation of 160 *ns* at 300 *K* was carried out initially.

After the stabilization of the protein structure, judged by the root mean square deviation (RMSD) of the backbone atom of the protein, the last structure of the simulation was re-solvated in a cubic box with 1.0 *nm* of solvent on all sides. All the conditions were kept the same as the previous simulation except a time step of 1 *fs* for numerical integration of the equations of motion was used. Coordinates were saved every 1 *fs*, and a 200 *ps* long trajectory at 300 *K* was similarly obtained for the analysis of water dynamics.

