## Supplementary Document for "An algorithm-based investigation reveals the differential dynamics of water inside protein cavity as a function of distance from its wall"

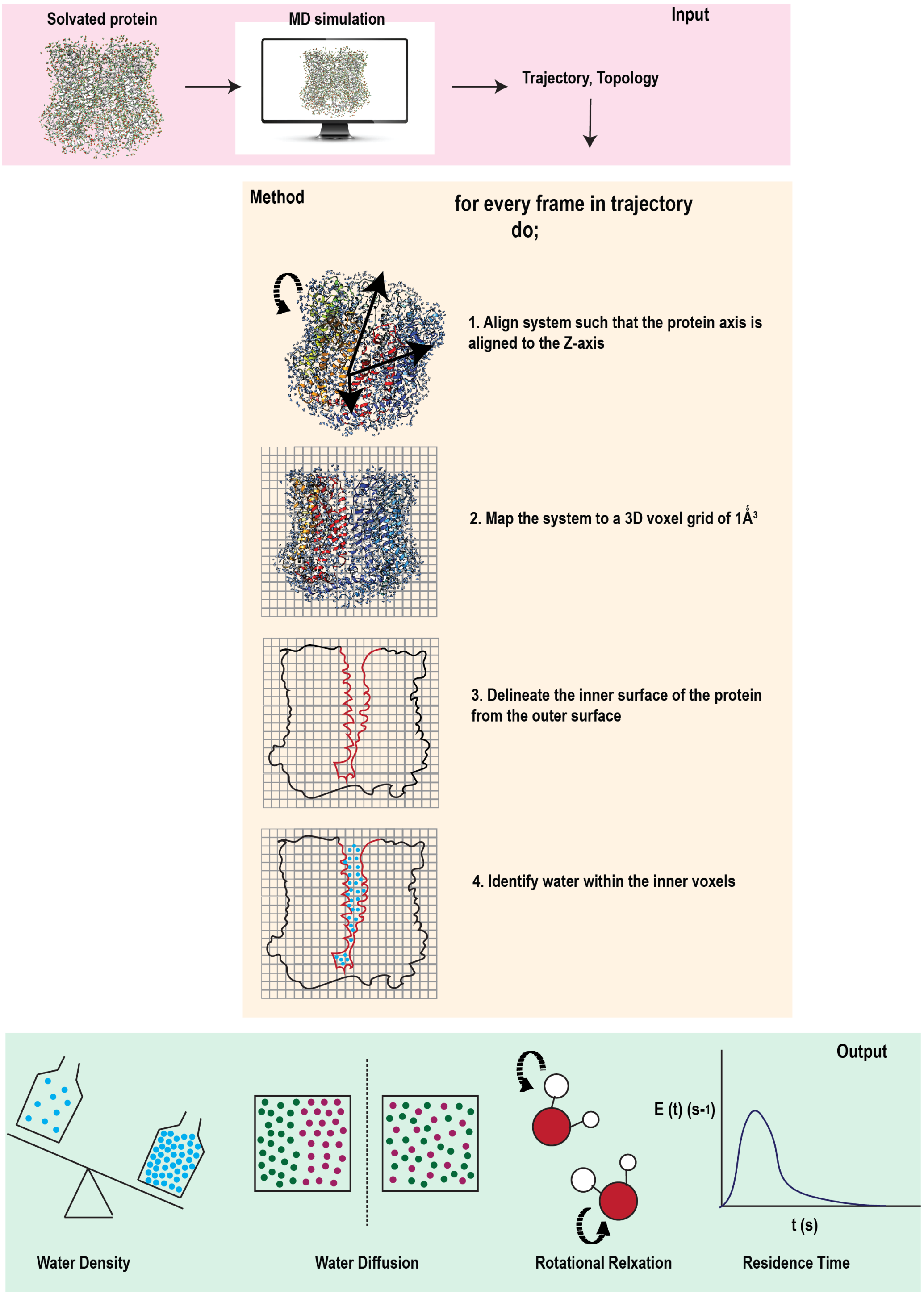


**Fig S1**: Schematic representation of the algorithm for the detection of water inside the inner cavity

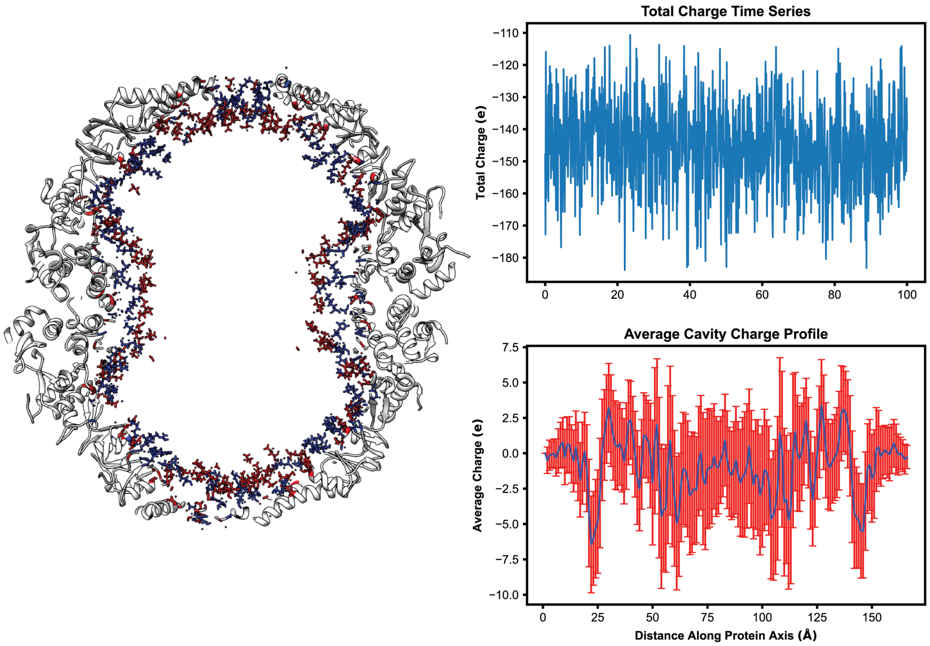


**Fig S2** : Charge profile of the inner cavity of Mm-Cpn (**PDB ID: 3LOS**). A thin cross-section of the cavity with negatively charged inner atoms (red) and positively charged inner atoms (blue) is depicted. The total charge of the cavity throughout the simulation (top right) and the average charge along the protein axis (bottom right) are also shown

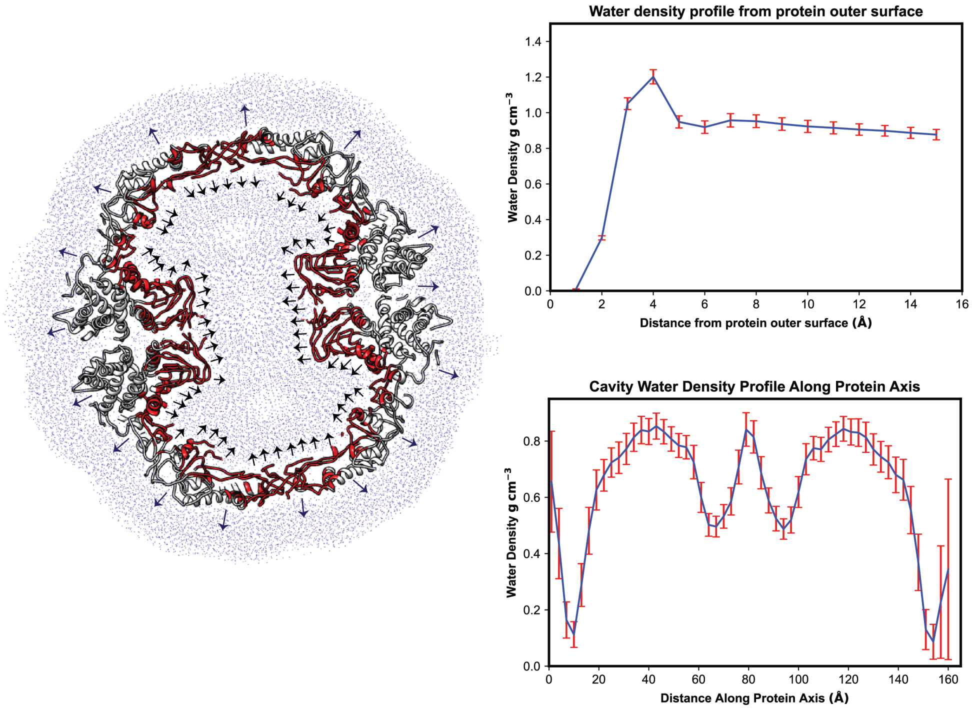


**Fig S3**: Schematic representation showing the direction for calculation of water density within the cavity of Mm-Cpn (**PDB ID: 3LOS**) and outside the protein (left). Density profile of water from the outer surface of the protein (top right). Water density profile in 1 $\mathring{A}$ steps along the cavity axis (bottom right). The thick line represents the mean density in $g\,cm^{-3}$, while error bars represent deviation from the mean density.

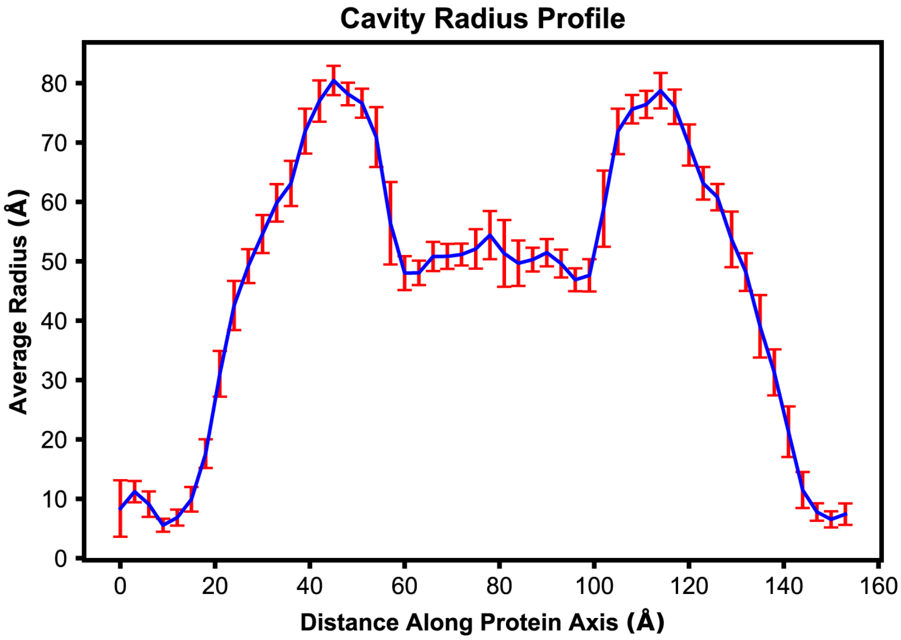


**Fig S4**: Dimensions of the cavity depicted by the radius of the inner cavity of Mm-Cpn (**PDB ID:3LOS**) along the protein cavity axis.

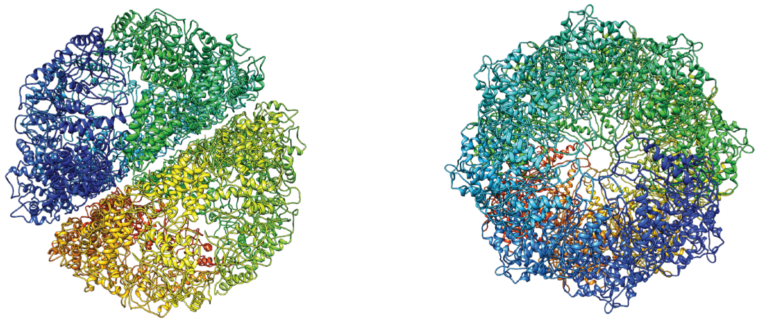

| System name | System size # of atoms | Protein size # of atoms | # of water atoms | Time |
|---|---|---|---|---|
| 3los | 698,652 | 126,176 | 572,124 | 160 ns |
| 3los-fs | 817,227 | 126,176 | 690,699 | 50 ps |

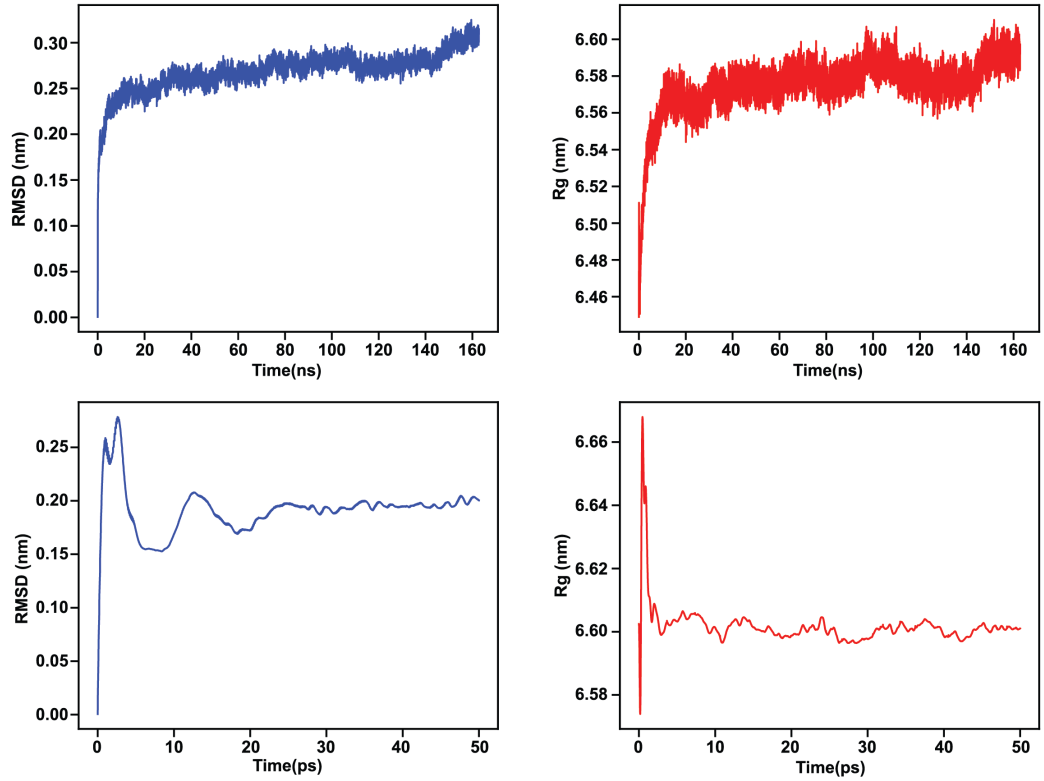


**Fig S5: MD simulation details for Mm-Cpn (PDB ID: 3LOS) used in the study** Simulation stabilized structure of Mm-Cpn is shown along with the root mean square deviation ($RMSD$) of the protein backbone atoms and radius of gyration ($R_g$) of the protein for both the simulations is shown.

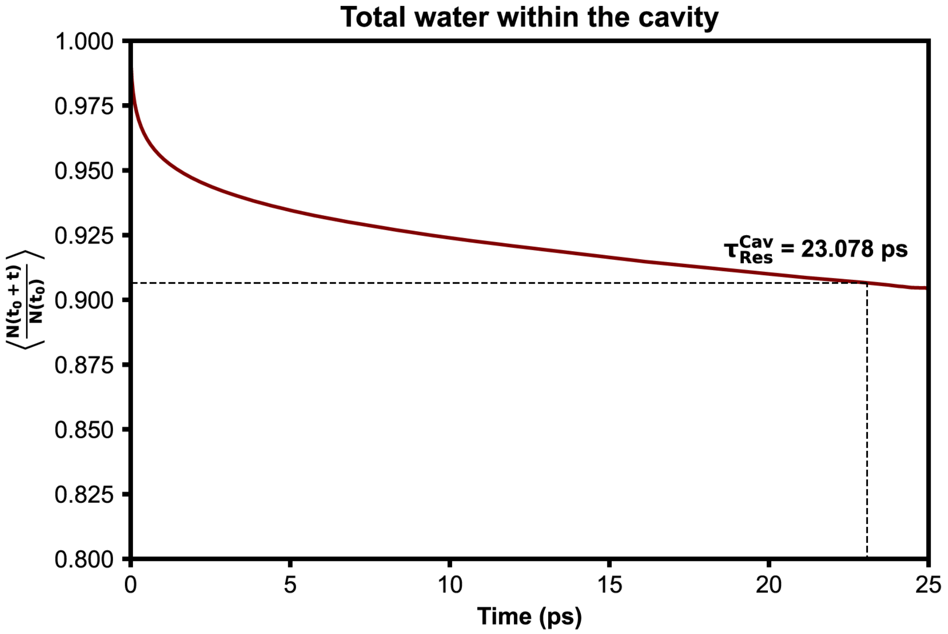


**Fig S6: Residence time of the total water within the cavity of Mm-Cpn (PDB ID: 3LOS) calculated using our method.**

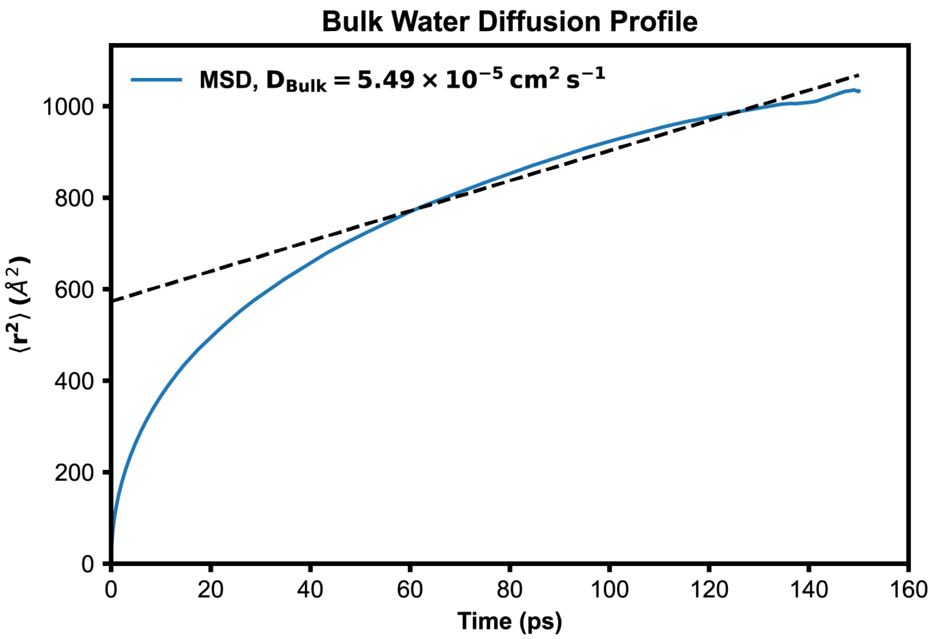


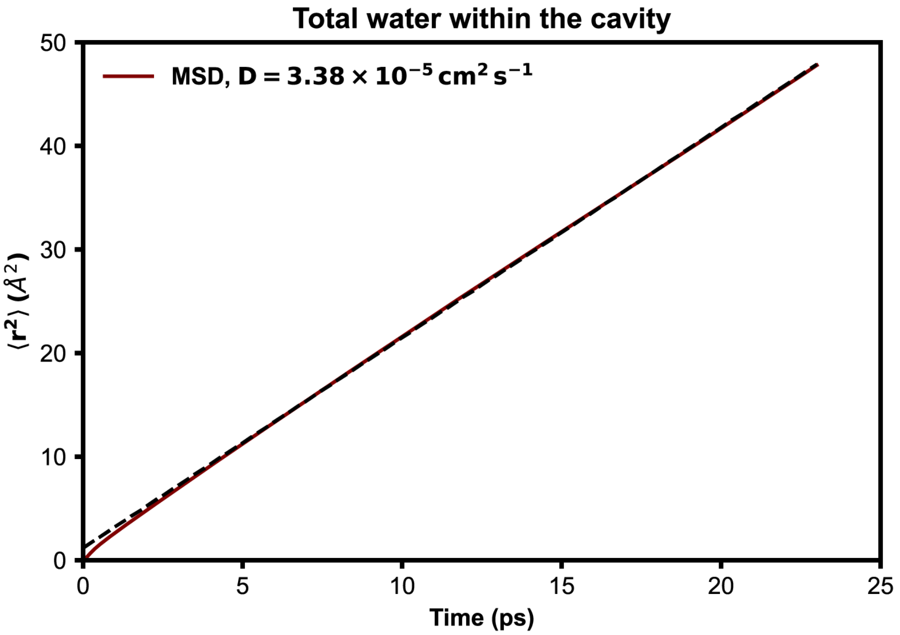


**Fig S7:** Diffusion profile of bulk water (calculated by simulating 4055 water molecules in a water box of dimension with density (top). Mean square diffusion profile of the total water within the cavity of Mm-Cpn (**PDB ID: 3LOS**) (bottom)

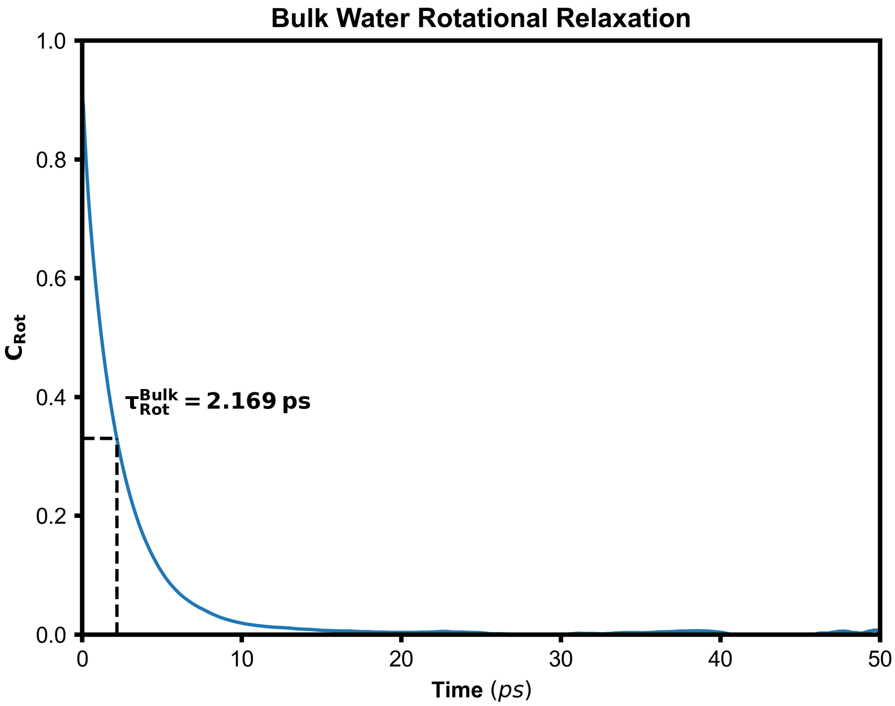


**Fig S8:** Rotational relaxation of bulk water calculated by simulating 4055 water molecules in a water box of dimension with density

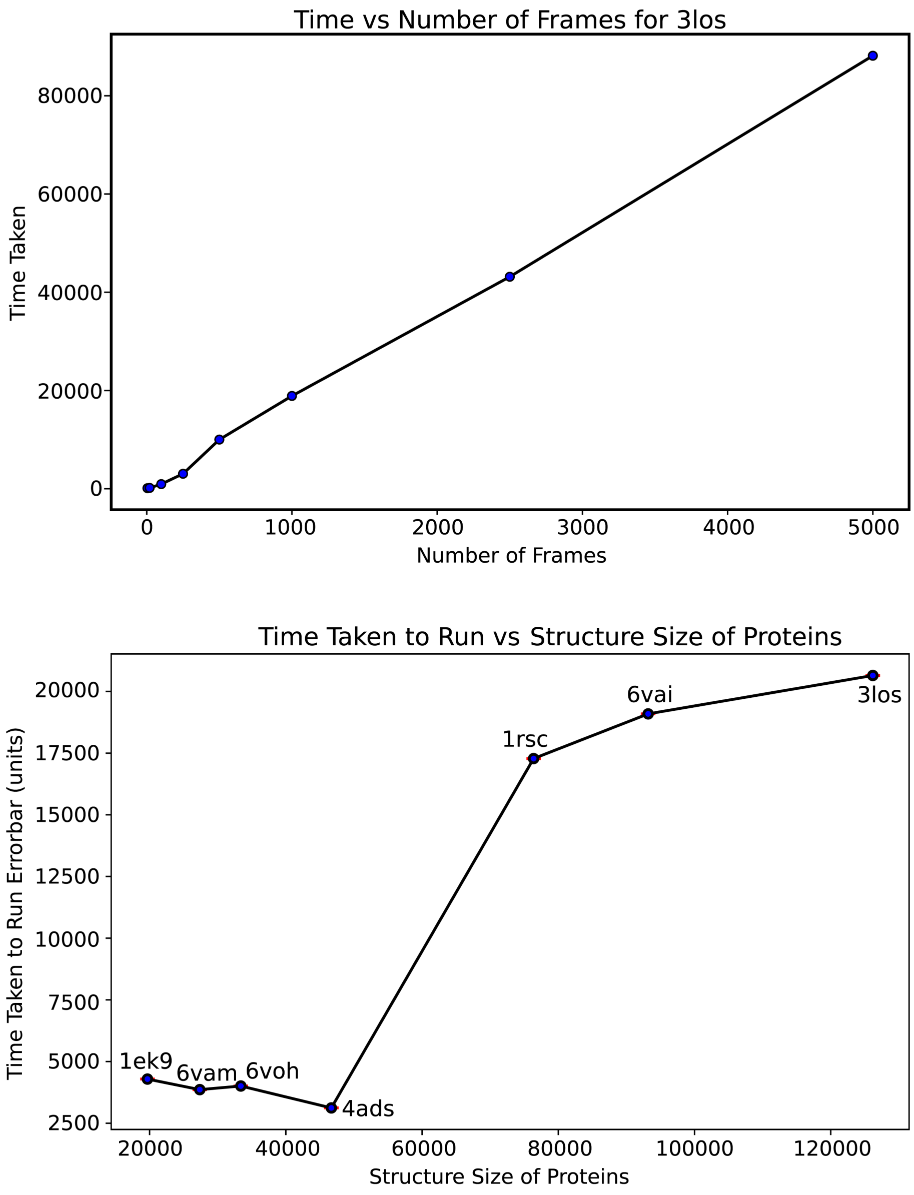


**Fig S9: Method optimization**. The time taken to identify cavity water increases linearly with the number of frames in the trajectory (top). The time taken to identify cavity water for 1000 frames for proteins varying in size (bottom).
